# Eyes on VR: Unpacking the Causal Chain Between Exposure, Reception, and Retention for Emotional Billboard Messages

**DOI:** 10.1101/2024.07.19.604208

**Authors:** Hee Jung Cho, Sue Lim, Monique Mitchell Turner, Gary Bente, Ralf Schmälzle

## Abstract

The causal chain from message exposure to reception to effects is widely accepted as the basic explanatory model for communication outcomes. Problematically, the chain’s links are often studied in isolation, leaving measurement gaps that compromise the ecological validity and practical utility of experimental research. Here we introduce a VR-based paradigm that encompasses a realistic message reception context, i.e., a simulated car ride on a highway flanked by billboards. We varied attentional message factors (emotional content) as well as contextual task distractions (trash-counting). VR-integrated eye trackers were used to capture participants’ incidental message exposure dependent on their actual gaze behavior. Consistent with our predictions, results show that 1. exposure gates all subsequent effects; 2. distraction impacts likelihood of exposure; 3. both the manipulation of emotional content and distraction affect retention. This comprehensive analysis of the exposure-reception-retention chain can be broadly applied to a variety of message reception contexts that will be discussed.

## 1. Introduction

Humans encounter a deluge of messages daily, but amidst continual opportunities for exposure, many messages are completely ignored, some receive close attention, and few are retained in memory (e.g., 1). For instance, consider the myriad billboards people pass by during routine commutes. Which ones remain in individuals’ memory? What factors contributed to retention of one over others? Does the memorability of remembered messages stem from heightened states of attention of the observer, or is it attributable to features inherent to the message itself, like their emotional content? Or is it the synergistic effect of both factors?

While encountering a multitude of visual messages, people are typically free to look at or ignore the messages. Exposure (2), the contact between the message and the recipient, depends on selective visual attention. Media effects hinge on whether individuals even look at a message, and those messages that are not seen cannot have any influence. However, despite the importance of exposure for communication, our knowledge of exposure in real-world contexts is grossly inadequate. Even after decades of empirical communication research, we neither have good estimates of how many messages a typical individual encounters on a normal day, nor what fraction thereof they attend to or ignore. Progress in screen-based analyses (3) can offer such estimates for computer-mediated messages, but a severe measurement gap persists for natural communication contexts in which messages are embedded in complex environments with competing attentional demands, and where message exposure is incidental and depending on peoples’ idiosyncratic behaviors. Moreover, while measuring exposure is crucial for understanding message effects, it is also only a necessary first step and ultimate message effects depend on further processing steps related to how we engage with messages, and how we store them in memory (4). This paper examines these fundamental questions for mass communication and media effects theories, with significant practical implications for health and political communication as well as advertising.

The paper is structured as follows. First, we introduce the causal chain from exposure, to reception, to retention, discussing how overt attention converts exposure opportunities (messages floating in one’s environment) into actual reception (messages that are looked at and inspected), and how paying attention to messages facilitates subsequent retention (memory). Second, we discuss how previous research is generally compatible with this model but suffers from key gaps regarding quantifying exposure in a rigorous manner while striking a balance between experimental control and realism. The current study uses VR to create a messaging context (a drive down a highway with billboard messages alongside), employs eye tracking to rigorously quantify the exposure-reception nexus, and manipulates both the presented billboards’ emotional content and the drivers’ attentional resources to demonstrate the hypothesized causal links among these theoretical variables.

### 1.1. Attention and the Causal Chain from Exposure, to Reception, to Retention

Attention is a fundamental bridging construct between mass communication (focusing on messages in the external information environment) and psychology (focusing on the encoding and processing of those messages inside the neurocognitive system; 5-7). Broadly defined, attention refers to the cognitive process of selectively focusing on a particular aspect of information (8, 9). However, rather than treating attention as a unitary concept, we can distinguish various subtypes based on the setting, task, or other characteristics (10).

In the context of visual information environments, such as billboards along a highway, a first kind of attention is overt visual attention (11). Overt attention refers to the fact that one can attend to (look at) aspects of the field of view, for instance by fixating on a billboard while averting gaze at least for a moment from the road. This kind of attention functions as a gatekeeper for all subsequent message effects. Understood this way, the act of attending overtly to a message links exposure to reception (5), which is what we turn to next. However, even when we overtly look at information, as in reading this sentence, we can process it in a more focused or more superficial manner (12, 13). Thus, even when exposure is sure, there is a secondary kind of attentional selectivity, that is how long or how deeply we engage with content (e.g., 14). There is evidence that links this kind of attention to memory (e.g., 15, 16).

Although the literature on visual attention is vast, two generalizations can be made regarding the modulators of attention: message characteristics and task demands. First, regarding the message characteristics, messages that are salient or conspicuous attract and sustain attention (17). Salience can be defined narrowly via features like brightness, contrast, or sudden onset; these lower-level attributes also explain a large share of eye movements and thus overt visual attention (18). However, above and beyond lower-level attributes, higher-level attributes like emotional content can also modulate attention. For example, pictures of cute babies, threatening or scary images, or explicit content all tend to capture and hold visual attention (19-21). Applying this to the driving context, highly emotional billboards might attract attention more powerfully than less emotional billboards, and there is evidence that drivers passing by accident sites exhibit ‘attentional rubbernecking’ effects driven by emotionality (e.g., 22). Extensive research on emotional messages shows a link between emotionality, attention, and memory (23, 24), although this is mainly demonstrated for very strong emotional content, less so for the more delicate touches of emotion we see in daily media messages.

Second, regarding task demands, it is well known that when attention is focused on one task, the performance on another task can suffer (e.g., 25, 26). For instance, when we are looking out for something in particular, we may fail to notice even very obvious and large objects (27). In typical experimental contexts, we can steer participants’ attention by instructing them to look out for and count particular target items (28). This can also be applied to the context of driving where one primarily focuses on driving, but other items may compete for attentional resources (29).

### 1.2. The Memory Trace: What Sticks after Exposure and Processing

Summarizing the above, we argue that the links of the causal chain go from exposure to reception and to retention (e.g., 4, 30). Attention (overt visual) is critical for both turning exposure opportunities (messages one could look at) into reception - making sure that they are encoded in the first place. However, attention also refers to how intensely people engage with messages during the post-exposure reception process thus affecting how they are encoded (e.g., 25, 31). Furthermore, we know that emotional messages may attract and hold attention better, and we know that distraction interferes with or depletes attention. All these variables should thus affect the chance of retention in a predictable manner: First, only exposed messages have a chance to make it into memory. However, not all messages we see, and process can be stored verbatim. Thus, more emotional messages should command more attention and generally facilitate memory formation. Finally, attentional distraction should reduce exposure likelihood and dampen subsequent processing, thus lowering the likelihood of remembering a message.

### 1.3. Confronting Complexities: The Challenge of Examining this Exposure-Reception-Retention Pathway

This theoretical chain from exposure to reception to effects is a logical and generally accepted explanatory model across communication. The processes described above (overt attention, selective attention, and memory encoding and retrieval) have been intensively studied in both cognitive psychology and neuroscience (32). Moreover, work on the processing of media messages is also compatible with this reasoning (25), and so is McGuire’s classic matrix of persuasion (4), or other models in mass communication and advertising research (13).

However, all prior work differs in important ways from the current experimental context (billboards along a highway), particularly regarding stimuli, tasks, and external validity: First laboratory studies in experimental cognitive psychology and neuroscience often bear little resemblance to the more natural phenomena they were designed to study, particularly not to everyday messaging contexts (e.g., 33). Second, even though prior work on mediated messages aligns well with the information processing view presented above, such work has typically focused only on temporal media messages (e.g., TV and radio spots; 34). Most critically, it has been done in laboratory tasks in which participants are force-exposed to messages. Thus, this work provides insights into post-exposure processing, but not in the exposure-reception link that requires studying peoples’ more active information search in realistic communication environments. In sum, if we are interested in research on exposure, then prior research provides only limited insight.

Research about message exposure does exist, but it is rather disconnected from the reception-focused research. Specifically, in mass communication and media effects research, exposure is a core theoretical concept (2, 35). Dozens of metrics exist that focus on audience size (how many are exposed to messages), reach, or frequency of messaging and effects of e.g., repeated exposure etc.. However, it is important to note that the majority of this work on exposure regards aggregate level exposure metrics, like reach and audience sizes, but not about whether a given individual looks at a message and how that influences the individual (36). In summary, despite the fundamental importance of the exposure-reception- retention chain, it seems surprising that only scarce and fragmented research exists that connect these links. To the extent that they are connected at all, research suffers from either a micro-macro divide or a divide between work that focuses on either message exposure (in the real world) or processing (in the lab).

### 1.4. Combining Virtual Reality and Eye Tracking to Examine the Exposure-Reception- Retention Chain

We believe that eye-tracking combined with virtual reality (VR) presents a promising innovation that propels theoretical progress by enabling communication researchers to unpack the exposure-reception-retention chain. First, because eye-tracking measures directly where people look and for how long their gaze stays engaged, it provides objective information about exposure and represents a widely accepted measure of visual attention (37).

Second, VR offers a promising way to overcome the limitations of laboratory studies, especially their limited realism. Specifically, as its name suggests, VR provides a way to create virtual, but realistic environments. This may sound tautological, but the implications become clear if one considers the challenge of measuring real-world exposure discussed above: Typically, we cannot objectively know whether a person looks at a billboard when driving down a highway because we neither have eye-tracking information, nor can we manipulate experimentally which messages appear along a highway. On the other hand, experimental research often suffers from limited generalizability, particularly when done in restricted laboratory contexts (e.g., 38). With VR, it becomes possible to overcome this bottleneck by creating realistic communication environments (e.g., a road with billboards, a mall, or a city; 39). Because recent VR-devices have integrated eye-tracking, it becomes possible to combine the potential of creating realistic communication environments in which users behave naturally with the benefits of measuring eye-tracking.

Finally, VR-based experimentation allows to isolate and manipulate theorized variables (like the influence of message emotionality or distraction) while controlling confounds. This is a key prerequisite for establishing a causal chain and achieving strong inference (40, 41).

Recent research has already made some progress towards these goals. Specifically, Bonneterre (42, 43) and Schmälzle et al. (44) both used the same core idea combining immersive VR with eye-tracking to zoom in on the exposure-reception link and study it under controlled but realistic conditions. The former authors studied the reception of tobacco-related messages in the city-environment and demonstrated that incidental exposure can be studied and linked to memory outcomes and smoking-related attitudes. The latter authors introduced a VRbillboard paradigm developed around a highway-driving scenario and demonstrated the key role of incidental exposure as well as how driver distraction makes it more or less likely. This study builds on this prior work to take it to the next level by manipulating multiple variables related to attention - message emotionality and distraction.

### 1.5. The Current Study and Hypotheses

The current study tests the above-described theoretical framework in which reception is the central link in the causal chain from exposure to memory, with attention serving as a key modulator of both the exposure-to-reception and the reception- to-memory linkages. Examining this framework in a causal-experimental fashion is enabled by an innovative experimental-ecological approach: Specifically, we build on previous work (44) that combined VR technology with integrated eye-tracking to create a message reception context in which people are free to attend to or ignore messages they encounter (driving down a highway with billboards) while allowing us to rigorously manipulate variables, capture eye-movements, and examine how these factors affect incidental message exposure and subsequent retention.

With this approach, we conceived the following manipulations to influence attention (see Figure 1): First, we manipulated the emotionality of the messages that participants encountered. Specifically, we created different versions of the same billboard message - varying only the level of emotionality (low vs. high) while keeping other visual and textual features highly consistent. Participants were unaware of these variations, as every person encountered a mix of emotional billboards during a virtual ride down a highway, but behind the scenes, one participant would encounter the low emotional version of one and the high emotional version of another billboard, whereas another participant would receive exactly the opposite patterns, thus controlling for confounds. Second, we instructed participants to either look out for and count trash items along the road (trash-counting condition) or freely drive down the road (free-viewing condition), assuming that this manipulation should markedly affect how much they would attend to billboards. During the ride, we then used the VR-integrated eye-tracking to unobtrusively measure whether they attended to each billboard (i.e. overt visual attention) and for how long they looked at it (i.e. intensity). Finally, once participants arrived at their virtual destination, we assessed their memory of the messages. With this setup, we are thus able to isolate the effects of billboard emotionality (i.e. low vs. high emotional visual content) and driving condition (i.e. trash-counting vs. free-viewing) on visual attention and memory, combining high levels of experimental control with ecological validity.

**Figure 1:**
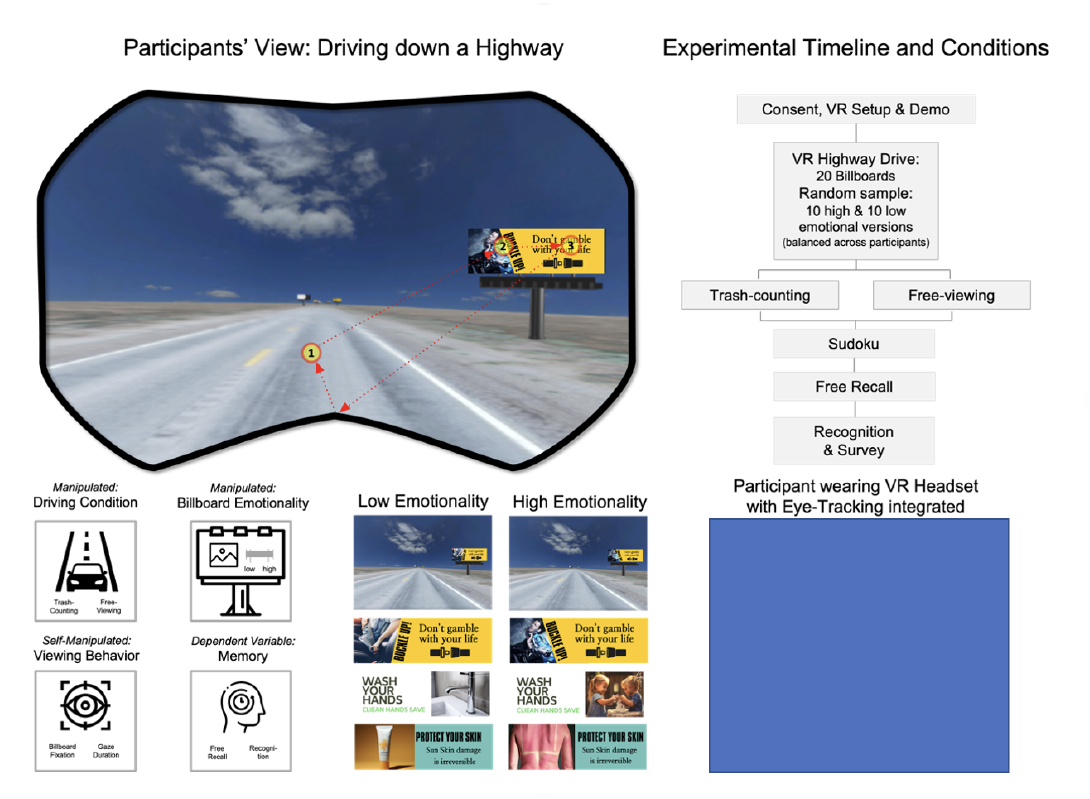
Study Overview: Experimental Setup and Manipulated Variables. Top left: Participant’s view of the photorealistic highway environment with billboards; Superimposed (not visible to participants) is the eye-tracking scan path, which is used to determine whether a billboard was looked at. Top right: Sequential diagram of study events and conditions. Bottom left: Independent and Dependent Variables. Middle Panel: Illustration of low and high emotional billboard versions. Bottom right: Lab setup: Participant wearing an HMD is engaged in driving along the virtual highway.

Based on the reasoning outlined above, the following hypotheses are proposed.

**H1**. Effect of Billboard Emotionality on Exposure and Reception: More emotional billboard messages will lead to more fixations (H1a) and longer gaze durations (H1b) compared to the less emotional billboard messages.

**H2**. Effect of Billboard Emotionality on Retention: More emotional billboard messages will be better recalled (H2a) and recognized (H2b) than less emotional messages.

**H3**. Effect of Driving Condition on Exposure and Reception: Individuals in the free-viewing driving condition will exhibit more fixations (H3a) and longer gaze durations (H3b) compared to those in trash-counting driving condition.

**H4**. Effect of Driving Condition on Retention: Individuals in the free-viewing driving condition will exhibit better memory performance (H4a measured via recall; H4b measured via recognition) than those in the trash-counting driving condition. We also conducted additional analyses to understand how viewing behavior intensity (i.e. fixation), driving condition (trash-counting vs. free-viewing) and message characteristics (low vs. high emotional content) influence memory.

## 2. Method

We provide code and data in a reproducibility package [link to data and scripts on OSF and Github will be included in the final manuscript]. In brief, participants wore a VR headset with integrated eye-tracking and drove down a photorealistic (virtual) highway along which billboard messages were placed (see Figure 1). Depending on the condition, they were instructed to count trash placed along the road or drive freely, and the displayed billboards were manipulated as described below. After the drive, participants’ incidental memory for the billboards was assessed via a free recall and recognition task. In the following, we report the specifics of the sample and procedures.

### 2.1. Participants

Forty participants (*m*_*age*_ = 20.2, *sd*_*age*_ = 1.5; 24 female) were recruited and received course credit. The study was approved by the local Institutional Review Board and all participants provided written informed consent. The sample size was set a priori to match the previous study’s sample (*N*=40) and is sufficient to detect expected strong effects of experimental manipulations on eye tracking and memory measures. Participants whose glasses did not fit under the VR HMD and with insufficient vision levels were immediately replaced, resulting in a final sample of 40 participants. Of these, 20 were randomly assigned to the trash-counting condition and 20 were assigned to the free-viewing condition (between-subjects).

### 2.2. Stimuli

We created visual billboards featuring various billboard-typical contents (see Figure 1). Specifically, out of the 20 billboards that every participant passed by while driving, 14 focused on health and risk topics, such as road safety, substance use, vaccination, and so forth; 6 of the 20 billboards focused on commercial topics, such as restaurants, coffee houses, hotels, or similar services. The billboards were designed using Canva.com and the Midjourney AI image generation tool to resemble typical billboard/outdoor advertising designs (i.e. text + images, see Figure 1 for an example and online repository [link to Github will be included in the final manuscript]). For every billboard, we created two versions, one featuring lower in image emotionality, the other higher in emotional content. In doing so, the text was always kept the same, but the images were varied in intensity of emotionality (low vs. high). To boost emotional salience of the images, the following elements were added: people, people with emotional expressions (e.g., startled due to an imminent accident), people in emotional situations (e.g., sad person after losing a loved one, person with a painfully red back due to sunburn, etc.). We confirmed that this manipulation of emotional salience was highly successful by having all participants perform a 2-alternative forced choice test (2AFC) after the study (dichotomous variable; 1 = low emotionality, 2 = high emotionality; see Table 1).

**Table 1:** Number and Percentage of Participants Who Correctly Identified Emotional Salience in the 2AFC Test.

| No. | Stimuli | *n* (percentage) |
| --- | --- | --- |
| 1 | Binge drinking | 38 (95) |
| 2 | Buckle up | 37 (92.5) |
| 3 | Diabetes | 28 (70) |
| 4 | HIV | 37 (92.5) |
| 5 | Text driving | 32 (80) |
| 6 | Drugged driving | 36 (90) |
| 7 | Smoking | 38 (95) |
| 8 | Sun protection | 39 (97.5) |
| 9 | Wash hands | 37 (92.5) |
| 10 | Vaccination | 35 (87.5) |
| 11 | Vaping | 38 (95) |
| 12 | Marijuana | 32 (80) |
| 13 | Technology addiction | 38 (95) |
| 14 | Healthy diet | 38 (95) |
| 15 | Brunch | 38 (95) |
| 16 | Burger | 37 (92.5) |
| 17 | Coffee | 38 (95) |
| 18 | Education donation | 39 (97.5) |
| 19 | Furniture | 35 (87.5) |
| 20 | Hotel | 34 (85) |

### 2.3. VR Environment and VR Device

The virtual environment that participants entered was a photorealistic version of Highway 50 in Nevada. This 3D-model was provided by the Nevada DOT on Sketchfab.com (link) and was equipped with additional details (e.g., a sunny and blue-sky dome with clouds, and empty soda cans on the highway for the trash-counting task).

The created billboard images were placed along this virtual highway on typical billboard stands. The order of the 20 billboard topics was fully randomized across viewers. Moreover, an elaborate assignment schema was devised so that pairs of participants received exact opposite emotionality versions (i.e., low vs. high) of the billboards, which were otherwise presented in the exact same order (e.g., sub001: #1: burger low emo, #2 hotel low emo, #3 drugs high emo; sub002: #1: burger high emo, #2 hotel high emo, #3 drugs low emo …). This mode of presentation ensures that all billboard versions are shown equally often and under conditions that are maximally comparable.

The VR headset was HP Reverb G2 Omnicept, which includes an embedded high-precision eye-tracker. Participants used the VR controller to drive forward in the virtual highway. There was no need for the participants to steer as the highway 50 model is perfectly straight.

### 2.4. Experimental Procedure and Conditions: Virtual Drive Down the Highway

After consent and VR preparation procedures were completed, participants first entered a demo environment to familiarize them with VR and how to use the controller to drive forward. Next, for the main session, they were instructed according to their assigned condition: Half of participants (*n*=20) were instructed to count the number of trash items on the highway (trash-counting condition; distraction condition), the other half were instructed to freely drive down until they reached the end (free-viewing condition, *n*=20). After the driving experience, which took about 5-7 minutes, participants were asked to work on Sudoku puzzles (2 minutes, serving to clear their working memory from the last billboards just passed), followed by a structured interview.

During this interview, the experimenter asked participants in the trash-counting condition how many trash items they counted and their general VR driving experiences. The interviewer then asked participants to list all billboards they could recall passing by (i.e., free recall). Finally, participants completed an online survey: This survey first asked them about VR experiences (spatial presence, occurrence of symptoms, and technology usability), followed by a visual recognition test of billboards. Specifically, for this recognition test, participants were shown all 20 billboards with both low and high emotional versions and 4 distractors. Then they were asked for each of the two versions of the billboard images whether they recognize seeing one of the versions while driving along the highway. If they answered yes to indicate they recognized seeing one of the two versions of the billboards, they were further asked to indicate their level of confidence. For this, the two billboard versions were shown at opposite ends of a bipolar matrix and participants indicated how confident they were having seen one or the other version (middle point indicating uncertainty). Last, participants viewed all billboard versions and selected the more emotional one for each alternative version (forced choice, manipulation test). Finally, participants were debriefed, and data were archived.

### 2.5. Data Processing, Analysis and Main Measures

This experimental setup yields the following objective data: First, from the VR-system’s output, we receive information about where participants were looking at, and particularly whether they fixated a given billboard while passing it (a dynamic/VR-based region of interest). In addition to assessing whether a billboard was fixated, we also measured for how long it was looked at in total (gaze duration) and how often it was looked at (in case of multiple fixations). By merging these viewing behaviors with the type of billboard that was displayed on a given position for a given participant (e.g., the billboard content as well as the low/high-emotional version), we can derive a list of which billboards and at which locations were viewed. Since billboard images were randomly allocated to specific billboard sign positions, a Python script was developed to reorganize the individual images based on a participant’s eye-tracking data (e.g., time 15s, billboard 1, drunk driving high emo.jpg, …). This facilitates subsequent data aggregation across participants and messages.

Second, from the interview and recognition survey, we can derive two metrics of message memory: free recall and recognition. These metrics are again captured at the individual level – i.e., whether participant X recalled banner Y, recognized banner Y, and which version of the billboard they saw. Thus, the central analytic dataset combines the following sources of information: 1) which billboard (e.g., smoking, texting and driving) and in which version (low vs. high emotionality) was displayed at which position (1,2, … 20) along the highway; 2) whether a given participant looked at (i.e., fixated) this billboard, how often this happened in the case of multiple re-fixations, and how long in total (gaze duration); 3) lastly, from the interview and the survey, we obtain measures of free recall and recognition, respectively (i.e., recalled/not recalled and recognized/not recognized). Overall, with 40 participants and 20 billboards, we thus obtain a data frame with 800 rows. 20 participants were in the trash-counting and 20 participants in the free-viewing condition, and each of the participants saw 10 low and 10 high emotional versions of the billboards.

In the analysis, we first conducted a stream of two separate repeated-measures ANOVAs to demonstrate the effects of our manipulations (driving condition: trash-counting vs. freeviewing and billboard emotionality: low vs. high) on the viewing behavior towards the billboards (fixations and gaze duration) and on memory for the billboards (free recall and recognition), respectively. Then we brought together the information about fixations (whether a billboard was actively looked at) and memory in a joint model together with the experimentally manipulated variables. Said differently, we can think of the viewing behavior metrics as another variable (representing the nexus where exposure turns into reception). However, contrary to typical laboratory studies where exposure is forced onto participants, our study let participants look freely (either completely freely or taxing their attention with a competing trash-counting task, which did, however, still leave them some choice). Thus, the variable viewing behavior varies from subject-to-subject based on their idiosyncratic viewing behavior. To statistically analyze these data, we specified a logistic (generalized) mixed effects model in which recall (or recognition respectively) formed the dependent variables and driving condition, billboard emotionality, and viewing behavior were the predictors of interest. Further, to account for potential differences between billboard messages and individual subjects, we specified those two variables as random effects and controlled for their varying intercepts (45-47).

## 3. Results

We used a VR-environment with an integrated eye-tracker to rigorously quantify message exposure and link it to message memory. Specifically, participants drove down a virtual highway along which billboards were placed, allowing us to manipulate billboard messages (less vs. more emotional variants) and tasks (trash-counting vs. free-viewing). Then we captured whether they fixated the billboards in passing (fixation vs. no fixation) as a ground-truth measurement of actual exposure. Finally, we measured message recall and message recognition.

### 3.1. Participants Subjective Experiences in VR

First, to demonstrate how participants experienced the drive, we examined their responses from verbal interviews conducted right after they came out of VR. Participants generally commented that they found the virtual highway drive to be realistic and captivating. This was further supported by the postexperimental survey data, which showed that participants reported a high level of spatial presence in the VR environment (*mean*_*spatial presence*_ = 3.61, *sd* = 0.71 on a scale of 1–5, with all items scoring above the midpoint; 48). Additionally, participants reported minimal symptoms such as dizziness, fatigue, or eyestrain (*mean*_*VR symptoms*_ = 1.42, *sd* = 0.36 on a scale of 1–4, with all items below the midpoint; 49).

### 3.2. Effects of Experimental Manipulations on Viewing Behavior and Memory

Next, we examined how the experimental manipulations (*Billboard Emotionality*: low vs. high and *Driving Condition:* trash-counting vs. free-viewing) impact participants’ *Viewing Behavior* (fixations on billboards and gaze duration) and *Memory* (free recall and recognition), assessed separately. Results for the effects of experimental manipulations on viewing behavior are illustrated in Figure 2, the results for the effects of experimental manipulations on memory are illustrated in Figure 3 and data are provided in Table 2 and Table 3.

**Table 2:**
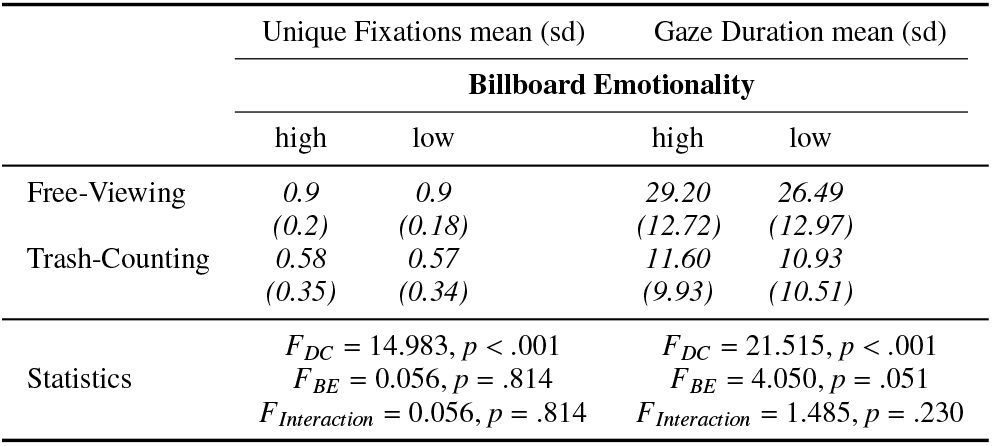
Means (SDs) and statistical results for effects of experimental manipulations (Billboard Emotionality/BE, Driving Condition/DC) on Viewing Behavior (Unique Fixations and Gaze Duration)

**Table 3:**
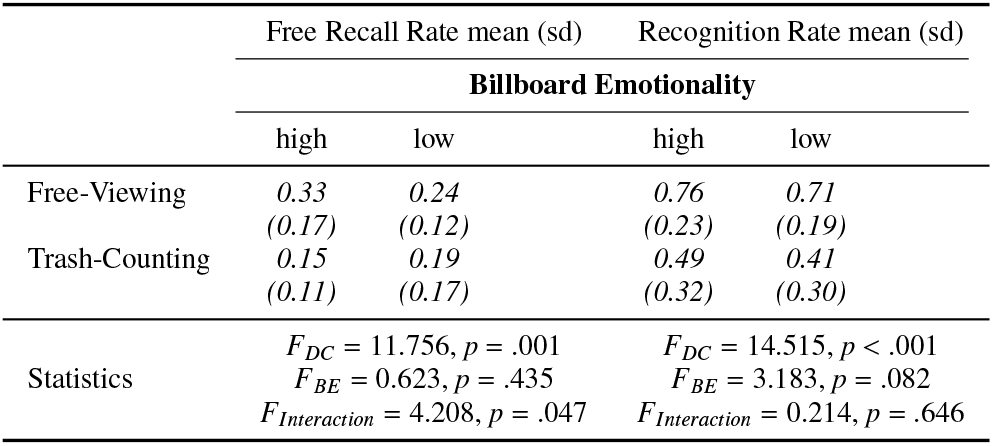
Means (SDs) and statistical results for effects of experimental manipulations (Driving Condition/DC, Billboard Emotionality/BE) on Memory (Free Recall and Recognition)

**Figure 2:**
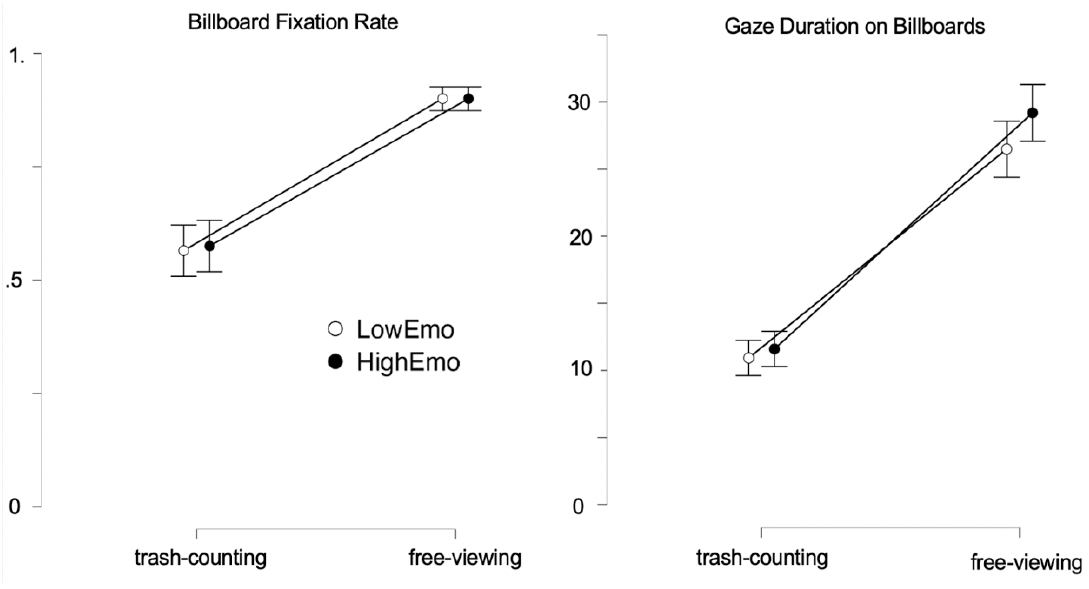
Effects of Billboard Emotionality and Driving Condition on Viewing Behavior. Left: Probability of fixating a billboard as a function of driving task (trash-counting vs. free-viewing) and billboard emotionality (low vs. high). As can be seen, in the free-viewing condition, participants are far more likely to fixate a billboard (on average 18/19 out of 20 billboards are fixated), whereas in the trash-counting condition, only about half of the billboards are fixated (i.e. looked at at least once). Right: Same analysis but for the gaze duration measure, i.e. sum of the total fixation duration across all billboards that were at least once fixated.

#### Effects on Viewing Behavior

As expected, participants in the ‘free-viewing’ *Driving Condition* were significantly more likely to fixate a billboard (ca. 90%) than participants in the ‘trash-counting’ driving condition (ca. 55%; *F*_*Driving Condition*_(1, 38) = 14.983, *p* < .001, see left panel in Figure 2). There was no main effect of *Billboard Emotionality* (*F*_*Billboard Emotionality*_(1, 38) = 0.056, *p* = .814) and the interaction effect between *Billboard Emotionality* (low vs. high) and *Driving Condition* (trash-counting vs. free-viewing) on fixation probability was not significant, *F*_*Driving Condition X Billboard Emotionality*_(1, 38) = 0.056, *p* = .814. Thus, the effect of the driving conditions on fixation probability was not significant regardless of the billboard’s emotionality.

A very similar pattern of results was obtained for the gaze duration as the dependent variable. As can be seen in the right panel in Figure 2, participants in the ‘free-viewing’ *Driving Condition* were not only more likely to look at a billboard, but they also looked longer at it if they did (*F*_*Driving Condition*_(1,38) = 21.515, *p* < .001). For the gaze duration measure, the main effect of *Billboard Emotionality* was marginally significant, *F*_*Billboard Emotionality*_(1, 38) = 4.050, *p* = .051, suggesting that the highly emotional billboards were looked at for a longer period. Again, there was no significant interaction effect, *F*_*Driving Condition X Billboard Emotionality*_(1, 38) = 1.485, *p* = .230. Thus, our results supported H3, and partially supported H1.

#### Effects on Memory Performance

Next, we examined whether participants’ memory (assessed via free recall and recognition, respectively) differed as a function of *Driving Condition* with their respective low (free-viewing) or high (trash-counting) attentional demands, and the *Billboard Emotionality* (less vs. more emotional content) factor (see Figure 3). Participants in the ‘free-viewing’ *Driving Condition* recalled significantly more billboards than participants who drove by the billboards while counting trash (*F*_*Driving Condition*_(1, 38) = 11.756, *p* = .001). The interaction effect between the recall rate of emotional messages (low vs. high) and the driving conditions (trash-counting vs. freeviewing) was significant (*F*_*Driving Condition X Billboard Emotionality*_(1, 38) = 4.208, *p* = .047). Participants in the freeviewing driving condition recalled more highly emotional billboards (*mean*_*recall rate: free-viewing, high emotionality*_ = 0.33, *sd* = 0.17) than low emotionality billboards(*mean*_*recall rate: free-viewing, low emotionality*_ = 0.24, *sd* = 0.12), whereas participants in the trash-counting driving condition recalled slightly more low emotional than high emotionality billboards(*mean*_*recall rate: trash-counting, high emotionality*_ = 0.15, *sd* = 0.11; *mean*_*recall rate: trash-counting, low emotionality*_ = 0.19, *sd* = 0.17). The main effect of *Billboard Emotionality* was not significant, *F*_*Billboard Emotionality*_(1, 38) = 0.623, *p* = .435.

**Figure 3:**
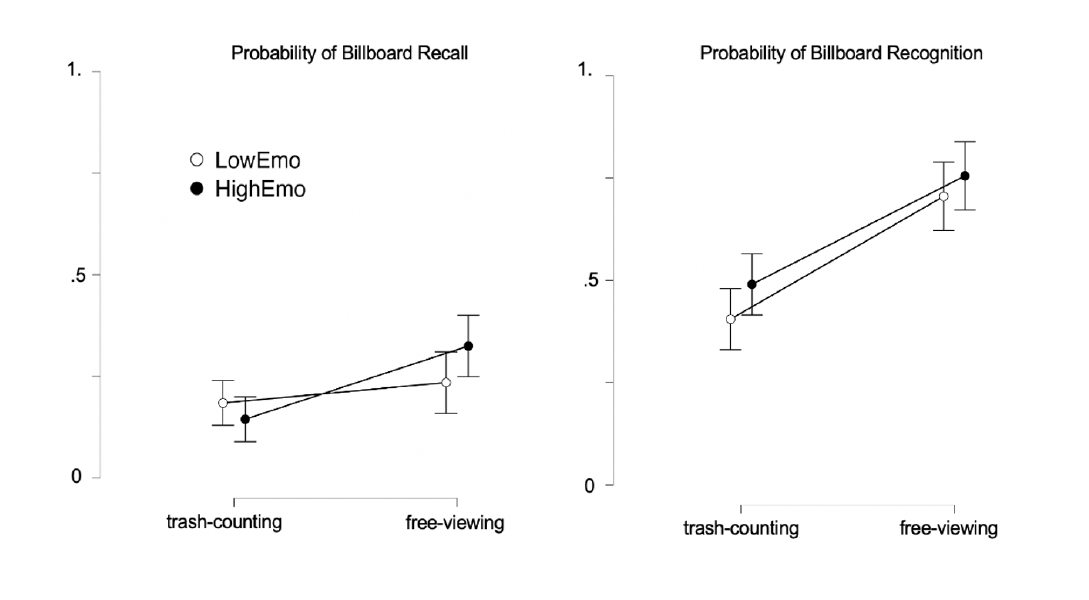
Effects of Billboard Emotionality and Driving Condition on Memory. Note. Left: Probability of Recall (i.e. freely mentioning a billboard after the virtual highway drive ended so that it could be identified). Right: Probability of Recognition, as measured in the post-experimental survey (note about guessing correction/signal detection analysis).

Performing the same analysis for the recognition memory revealed a highly significant main effect of *Driving Condition* (*F*_*Driving Condition*_(1, 38) = 14.515, *p* < .001): Participants in the free-viewing condition recognized significantly more billboards than participants who drove by the billboards while counting trash. The main effect of *Billboard Emotionality* was not significant, although a trend was seen (*F*_*Billboard Emotionality*_(1, 38) = 3.183, *p* = .082) for more emotional billboards to be recognized more often. For recognition memory, there was no interaction effect between *Driving Condition* and *Billboard Emotionality* (*F*_*Driving Condition X Billboard Emotionality*_(1, 38) = 0.214, *p* = .646).

### 3.3. Examining Message Memory as a Function of Viewing Behavior (Exposure), Driving Condition (Attentional Resources), and Billboard Emotionality (Attention-Attracting Image Characteristics)

#### Strong effect of Viewing Behavior on Memory

Having examined how the experimental factors *Driving Condition* and *Billboard Emotionality* impact participants’ *Viewing Behavior* and *Memory*, we next moved on to consider how *Driving Condition, Billboard Emotionality* and *Viewing Behavior* jointly impact *Memory* (see Figure 4). First, we confirmed that participants’ individual *Viewing Behavior* strongly impacts *Memory*. To this end, we specified a generalized linear mixed model (GLMM) to test the effects of all experimentally manipulated factors (*Driving Condition* and *Billboard Emotionality*) as well as the self-manipulated factor *Viewing Behavior* (fixation, whether a participants looked at a given billboard or not) on memory outcomes. For message recall as the outcome, a significant and dominant effect of *Viewing Behavior (fixation vs. no fixation on a billboard message)* confirmed that whether a billboard was fixated (looked at) or not strongly affects the probability of recall, χ^2^_*Viewing Behavior*_ = 43.430, *p* < .001. Likewise, conducting the same analysis for message recognition as the outcome, a significant dominant effect of fixation on recognition was observed, (χ^2^ _*Viewing Behavior*_ = 14.413, *p* < .001. Additionally, for recognition (but not for recall), the effect of *Driving Condition* (free-viewing vs. trash-counting) was also significant, χ^2^_*Driving Condition*_ (*df* = 1) = 10.953, *p* < .001. The results are illustrated in Figure 4, and they are consistent with the proposed causal influence of *actual exposure* as the dominant mediating link between message content and message effects.

**Figure 4:**
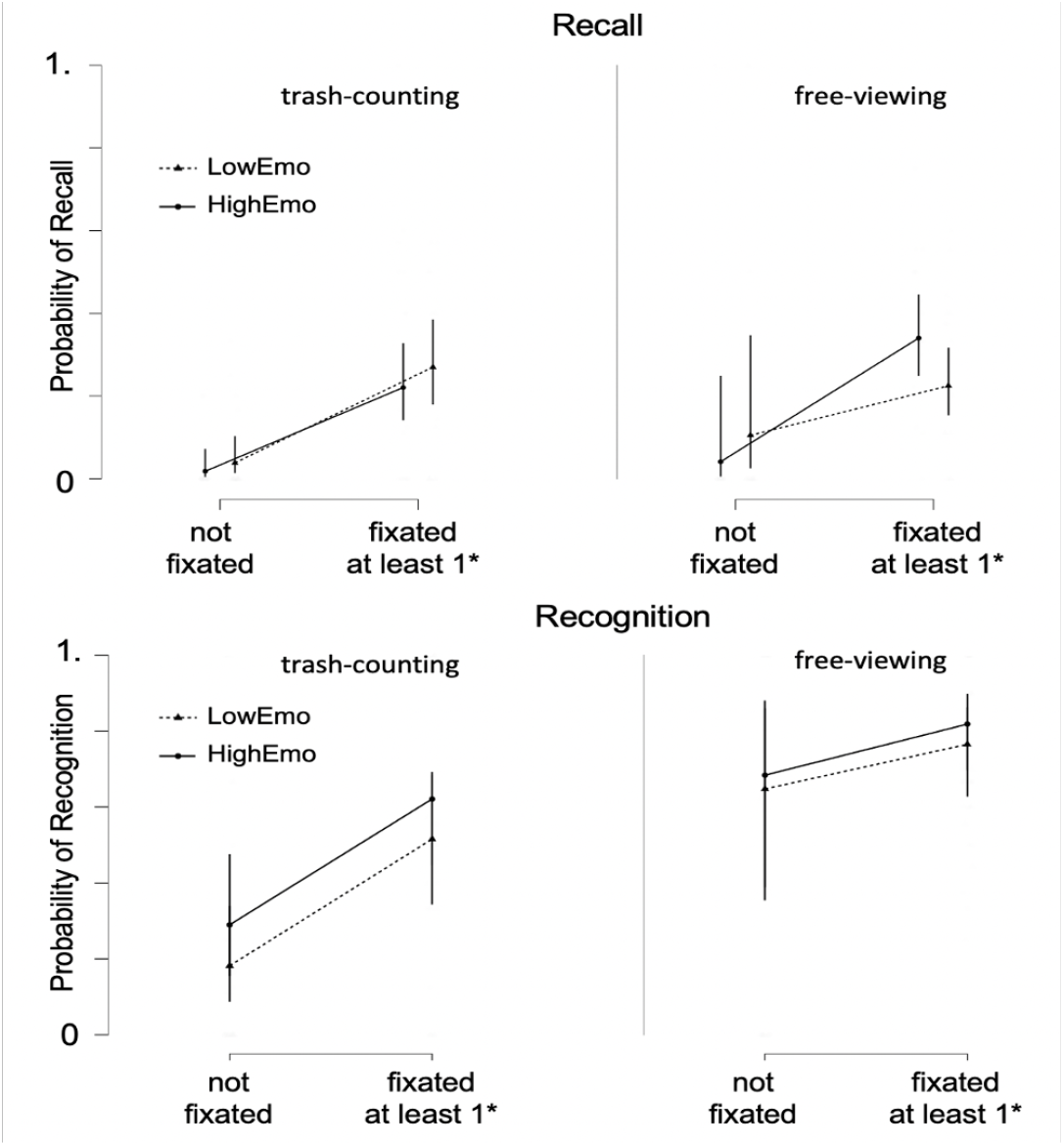
Joint model of experimentally manipulated variables (Driving Condition/DC and Billboard Emotionality/BE) and subject-determined behavioral manipulation (Viewing Behavior, i.e. whether a billboard was fixated or not) on memory outcomes. Top row: Results for free recall (left: trash-counting; right: free-viewing). Bottom row: Results for recognition memory.

#### Memory for Messages that were Looked At (exposed/actually seen) as a Function of Driving Condition and Billboard Emotionality

Having demonstrated that the self-determined *Viewing Behavior* strongly affects memory (both measured via recall or recognition), we zoomed in on only those billboards that were looked at, i.e. the ones for which we can objectively claim that participants were exposed to them. A generalized linear mixed model (GLMM) for the recall data from all looked-at billboards revealed a statistically significant interaction effect between *Driving Condition* (trash-counting vs. free-viewing) and the *Billboard Emotionality* (low vs. high), χ^2^_*Driving Condition X Billboard Emotionality*_ = 4.522, *p* = .033. This suggests that the impact of the billboards’ emotionality on recall rate varies depending on the viewing conditions. Specifically, for participants who were instructed to count trash, the recall rate did not differ much between the high and low emotionality billboards, but in the free-viewing driving condition (where participants resources were not taxed by counting trash), the recall rate was higher for high emotionality billboards compared to the low emotionality billboards.

Conducting the same analyses for the recognition memory revealed a significant main effect of *Driving Condition* (higher memory in the free-viewing condition, χ^2^_*Driving Condition*_ = 5.361, *p* = .021) and a marginally significant main effect of *Billboard Emotionality* (higher memory for more emotional messages, χ^2^_*Billboard Emotionality*_ = 3.645, *p* = .056).

### 3.4. Additional Analysis of Dose-Response-Relationships

*Does more intense viewing lead to better memory?* The analyses presented in the previous section are based on a dichotomous conceptualization of exposure, defined as whether participants did or did not look at a given billboard at all. However, a more nuanced view can be provided by a gradual analysis that focuses on dose-response relationships. To this end, we went back to the original data and reanalyzed for every participant and every billboard how often (fixation count) they looked at a billboard. Specifically, we split every participant’s fixation data to form three bins: billboards that were never looked at, billboards that were looked at some (but less than that participant’s median number of fixations) or looked at a lot (more than median number of fixations). For each of those three classes of billboards, we analyzed the subsequent memory performance. The results are shown in Figure 6 and revealed a very clear picture: For both, recall and recognition memory, significant main effects of viewing behavior intensity and driving condition were qualified by a significant interaction. Although even in the trash-counting driving condition more fixated billboards tended to be better memorized, these effects were more strongly pronounced in the free-viewing driving condition (Recall: *F*_*Viewing Behavior Intensity*_(2, 76) = 65.265, *p* < .001; *F*_*Driving Condition*_(1, 38) = 11.756, *p* = .001, *F*_*Viewing Behavior Intensity * Driving Condition*_(2, 76) = 5.499, *p* = .006; Recognition: *F*_*Viewing Behavior Intensity*_(2, 76) = 30.097, *p* < .001; *F*_*Driving Condition*_(1, 38) = 14.515, *p* < .001, *F*_*Viewing Behavior Intensity * Driving Condition*_(2, 76) = 4.975, *p* = .009). These data confirm dose-response relationships, which are an important topic in health communication at the aggregate level (50), but are demonstrated to matter here even at the level of micro-level reception data. Put simply, the more a billboard is inspected, the better the memory.

## 4. Discussion

This study examined the causal link between exposure, reception, and retention within a controlled experimental setting, and specifically the influence of attention-manipulating factors message emotionality and driver distraction (i.e., trash-counting (search task) vs. free-viewing (free inspection) on exposure and retention. Our results confirm that exposure determines all subsequent effects, that distraction impacts likelihood of exposure, and that both, the manipulation of distraction and emotional content, affect retention. Below we discuss these results and their theoretical significance, including how the current approach advances our mission to reveal the mechanisms from messages in the environment to their effects on audiences.

First, we observed strong effects among driving conditions, fixations, and memory. The participants in the free-viewing condition (i.e., non-divided attention) were about 1.5 times more likely to fixate on a billboard and look at it for a longer period (gaze duration) than those in trash-counting condition (i.e., divided attention, see Figure 2). As we predicted, the participants in free-viewing condition recalled and recognized more billboards (see Figure 3). These results align with existing studies about the link between attention and memory (e.g., 16, 33, 51). Thus, our results show that actual exposure (i.e., “fixating on the billboard”) explains variance in message memory (52), and they underscore the importance of studying the exposurereception-retention nexus in an integrated manner (44).

Next, we turn to the influence of message emotionality on gaze behavior (fixation likelihood and gaze duration) and memory. As can be seen in Figure 4 and contrary to our predictions message emotionality did not strongly affect whether or not people fixated on the message. However, high emotionality messages were looked at for a longer period compared to the low emotionality messages (marginal statistical significance), supporting H1b. Interestingly though, when zooming in only on the messages that were fixated (see Figure 5), we found that the effect of message emotionality on memory varied by the driving condition (i.e., available attention) and the type of memory. Participants in the free-viewing condition recalled slightly more high emotional messages compared to the low emotional messages.

**Figure 5:**
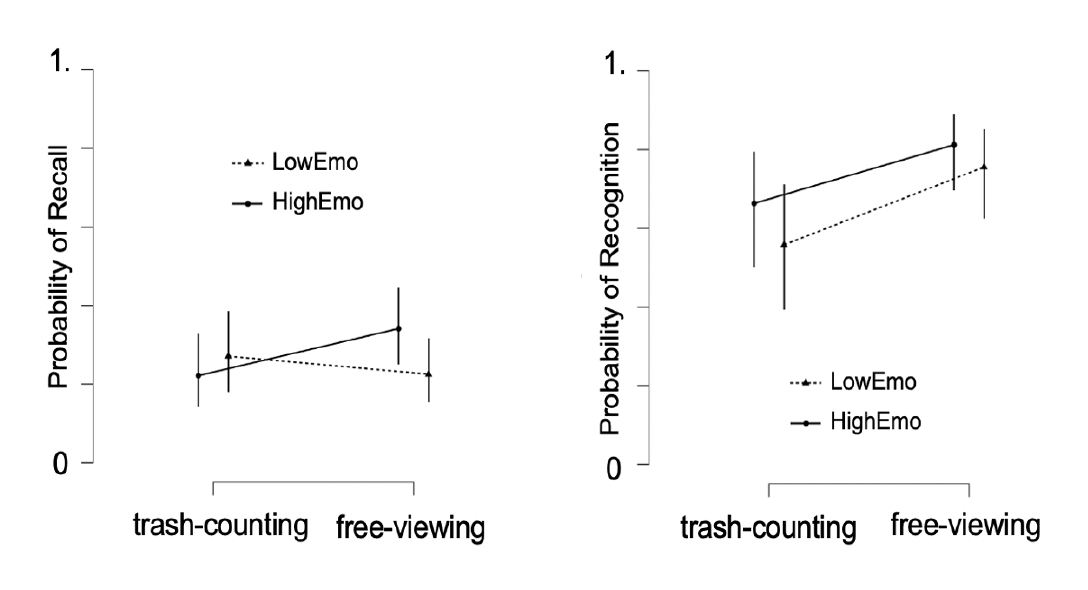
Zooming in on Effects of Billboard Emotionality (BE) and Driving Condition (DC) on Memory for Fixated Billboards Only (i.e. certain exposure) Note. Left panel: For the recall memory measure, a significant interaction is observed. Particularly during the free-viewing condition, highly emotional messages are remembered best. Right Panel: For the recognition memory measure, two main effects emerge. Free-viewing leads to better memory and more emotional messages are more often recognized.

**Figure 6:**
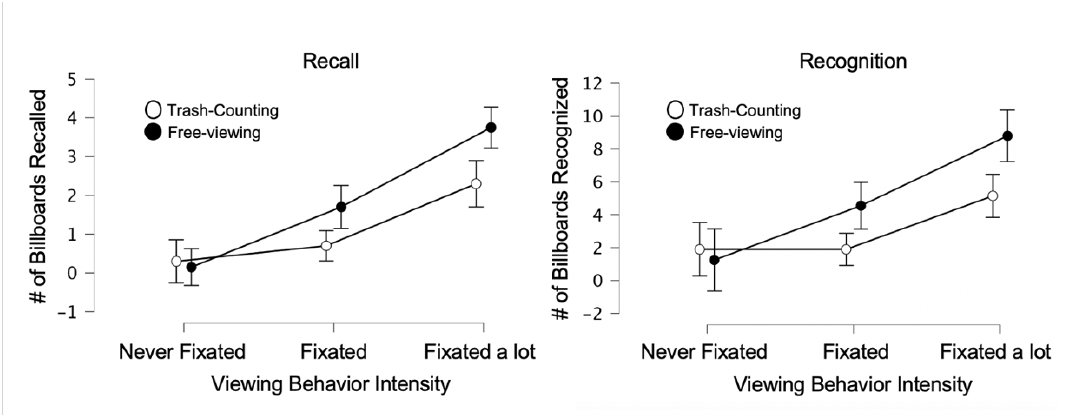
Results from Additional Analyses of Dose-Response Relationships

Taken together, our two manipulations impacted *whether* and *how* participants looked at and remembered the messages. The distraction manipulation had very strong main effects on viewing behavior (fixation rate and duration) and memory (recall and recognition). The effects of billboard emotionality were not as strong, and the most conspicuous effect was for fixated billboards (for which we can be sure that exposure happened) the recognition memory revealed a main effect of billboard emotionality (higher emotionality billboards being recognized better); this effect was similarly expressed across both driving condition groups, demonstrating consistency. However, when memory was assessed via free recall, which is more difficult as it requires the active retrieval of a memory trace, the pattern of results was different: Under free-viewing conditions, i.e. when less attention was consumed by the competing task of trash-counting, we observe higher recall of more emotional messages. But in the trash-counting condition, which required deploying attention on the road, the more emotional billboard versions were not more successful in attracting attention or boosting memory. One potential reason behind the observed differences in driving conditions; trash-counting vs. free-viewing may be discussed in relation to the distinct cognitive demands each impose on individuals. The introduction of trash-counting task is not merely a matter of adding visual distractions but rather it fundamentally alters the participants’ mindset by shifting from a free inspection approach to a more targeted search task which engages participants in a continuous alertness preparing them to respond to an anticipated task (c.f. 53). This imposes the cognitive load which constrains the available cognitive resources leaving less capacities for ones to deeply process the stimulus that differ in terms of its emotionality (low vs. high). Therefore, the task types may have influenced the cognitive resource allocation; the high cognitive load during the trash-counting task might have suppressed the salience of billboard emotionality.

Overall, by using VR and an integrated eye-tracker in this billboard paradigm, we were able to manipulate participants’ attention, unobtrusively capturing their viewing behavior in space and time, and linking this information to memory outcomes.

### 4.1. Implication 1: Measuring Actual Exposure

The theoretical significance of our results lies in the clear, simple, and objective insights they offer into the exposurereception-retention nexus. Specifically, the way in which exposure was measured in prior research leaves a lot to be desired: Measuring exposure at an aggregate level is different from measuring it in a given individual (2, 54); measuring exposure via self-report is potentially subject to recall bias (55); and last, laboratory studies can examine forced exposure, but not the kind of incidental exposure that matters in real life (56). Thus, it is safe to say that the theoretical phenomenon in question actual exposure in ecological settings has barely been measured. This poses a significant theoretical challenge insofar as exposure has been termed the foundation of all message effects (e.g., 35). In this sense, the current study makes not only a methodological contribution, but provides a great example of the old adage that there is nothing so theoretical as a good method (57).

### 4.2. Implication 2: Unpacking the Mechanisms of Message Emotionality

Beyond rigorously quantifying actual exposure, the current study manipulated the messages’ emotional content. The underlying reasoning was that there is a logical chain from message content (e.g., a billboard displaying a picture of road accident with a textual warning to avoid texting and driving) to exposure and reception (the driver passing the billboard and looking at it), and on to retention (the driver recalling or recognizing this message; 58). The goal of communication message design is to manipulate specific message characteristics to affect this chain from increasing the likelihood of exposure to facilitating sustained attentional engagement, to boosting memory encoding (e.g., 59, 60). For instance, by increasing the physical image salience (e.g., via contrast or flashing lights around the billboard), one can attract gaze, and the underlying perceptual and cognitive mechanisms are fairly well understood (61). Beyond image salience, however, most researchers are more interested in higher-level message characteristics, like the effects of emotional appeals, framing, or narratives (60).

Here, we focused on the emotional visual content of the messages, creating two well-matched versions of each billboard (one more, the other less emotional; cf. 62). This also aligns with decades of communication research, particularly the seminal work on the effects of fear appeals (63, 64), although we note that we did not manipulate fear specifically (most of our high-emotion billboards depicted high-arousing negative consequences, like accidents or sunburn, which would be considered a threat appeal). The assumption was that by making the visual billboard content more emotional, they would be more likely to be looked at, more intensely inspected, and ultimately more likely to be stored (65). Indeed, neuroscientific research on affective vision strongly supports that emotional images are prioritized and amplified across processing stages from early vision to memory formation (19, 66).

However, the current results only partially supported this. Although overall, more emotional images were somewhat better remembered, the effect of this manipulation were much weaker than the effects of the driving condition (distraction). Likewise, emotional images were not vastly more likely to be fixated, but once they were fixated, people indeed tended to look at them more often. In retrospect, these results make sense given that our emotionality manipulation was fairly weak. Specifically, research on emotional images tends to be done with very strong imagery, like pictures of strong mutilations and erotica (19), whereas our emotional content manipulations spanned across a more moderate range of emotionality (i.e. only depicting a startled driver, but not a blood-splattered car-wreck as in a full-blown fear/threat appeal). With this in mind, the current results seem reasonable and demonstrate that despite a fairly weak manipulation of emotional image content, some processing advantage is detectable (and our post-experimental forced choice task confirms that participants were able to detect the more emotional billboards). Although the effect size of this manipulation is moderate and barely significant with a sample of 40 participants, these effects could still matter practically if we consider that typical roadside billboards are passed by thousands of passengers every day (67).

### 4.3. Implication 3: Opportunities for Examining Message Effects in Real and Virtual Environments

Going forward, future work should expand the range of message characteristics and study the effects of those manipulations. As discussed above, it would seem possible to increase the strength of our emotion manipulation and doing so should lead to stronger effects. Alternatively, given that much emotion research also examines discrete emotions beyond the arousal/intensity axis, one could specifically zoom in on e.g. guilt appeals, humor, or other specific emotions (62). The messages with their own emotional properties, engage the viewer from the moment of the visual contact (i.e., catches the viewers’ eyes) through to cognitive appraisal (i.e., processes the content assessing/perceiving the emotional significance). Based on the cognitive appraisal, a decision is made whether the message warrants further attention (c.f., 68; Cognitive Functional Model). But the paradigm offers a lot of flexibility for additional manipulations. For instance, we did only manipulate the image content, but kept the billboards’ textual content perfectly controlled. It would seem easy to also manipulate text content, adding e.g. framing manipulations, short narratives, testimonials, or other kinds of manipulations. Given that recent work by O’Keefe and Hoeken (69) questioned the effects of numerous message manipulations, it appears promising to pit various manipulations against each other, creating a sort of “persuasion competition” along the virtual roadside.

We also note that much of eye-tracking based message effects research is limited insofar as it uses stationary eye trackers (70-72). Stationary eye-trackers offer valid insights into scanpaths on websites, but it limits the messaging contexts to screen-based-paradigms. However, if the goal is to understand how people look for and react to messages in more naturalistic information environments, a different approach is needed (38, 56, 73). In case of billboard advertising, for instance, people navigate freely through space, their eyes wander and constantly select information, and the relevant target objects (in this case billboards) change size as people approach and then pass them. The strength of the VR- and eye-tracking paradigm presented here is that it cannot only cope with these complexities, but that it also offers great flexibility in terms of potential messaging contexts that could be studied. Indeed, the same basic argument can be made for other kinds of outdoor advertising, like inner cities, malls, etc., but it could also be made for e.g. finding signs in hospitals, hallways etc. (e.g. 42, 43). As long as relevant environment and message features can be isolated, we can now manipulate them and examine how VR-immersed users behave when experiencing these carefully crafted, but realistic communication ecosystems (39).

The current paradigm also has broad applied implications, particularly regarding messaging and advertising in natural and future metaverse communication environments. In addition to the reliance on objective measures (overt attention captured via eye-tracking), a core strength of the VR-based approach taken here is its near-infinite flexibility regarding the types of contexts one could study: The current paradigm was situated in a highway and billboard advertising context, but it is easy to see how this can be transferred to e.g. an urban environment, a railway station, or any of outdoor advertising. But even in the highway-driving context alone, we see several practical applications, like providing legal evidence about impact of roadside advertising, empirical billboard efficacy measurement, or simulations for billboard construction planning. Last, and perhaps most importantly, we turn to the upcoming metaverse environment and its implications for this type or empirical communication research. In the case of the Metaverse, the VR-based environment will no longer be a model for the real world, but rather VR will be the context in which people spend time and encounter messages. This means that the current approach can be directly applied to study user engagement with messages. Given the enormous importance of digital user metrics (clickthrough rates, page impressions) had on the internet and its leading platforms (Facebook, Twitter, YouTube, Instagram), it will be important for communication researchers to 1. study this new messaging context, 2. use the new opportunities to advance theoretical communication science, and 3. also discuss the ethical and societal implications of this development (e.g. 74).

### 4.4. Strengths and Limitations

The strengths of our study include the following. First, we controlled how much attention people have available to allocate to the messages (trash-counting vs. free-viewing) and what types of messages they can be exposed to (low emotionality vs. high emotionality). Compared to other studies that are more focused on naturalistic environments (where there is eye-tracking or potential for message exposure but no control over the messages) or strictly lab-based work where people are forced to read/see every message, our study holds a middle ground. In addition, we also used advanced statistical analyses including generalized linear mixed models to take into consideration of individual differences in participants and messages in terms of outcomes we were interested in. Finally, we added innovative methodological (e.g., used AI to generate messages) and theoretical factors (e.g., added emotionality) compared to existing studies.

However, there are still limitations to this study. First, we purposefully controlled for the variance in the message manipulation because we did not want to cause any unintentional emotional harm to the participants. This could have backfired, leading to small or statistically insignificant effects of emotionality. Future studies can add more extreme emotionality manipulations and see if they lead to statistically significant effects. In addition, while VR and eye-tracking methods have advanced far, more objective measurements such as neural activity recordings such as the EEG could further add to our understanding of how attention, exposure, and message factors interact to influence memory. In particular, eye-tracking is a perfect measure of overt attention and can completely ascertain if a message was looked at. However, for the subsequent processing of that message, we can gain some insight from eye-tracking (e.g., how long people look – or we could even look at the pupil dilation as a measure of arousal), but it is clear that further measures are needed to fully unpack how the processing of messages is transformed into retention. Additionally, while the current study focused on trash-counting as the distracted condition (i.e. task type), the future research could benefit from incorporating real-life distractors such as unexpectedly and spontaneously appearing objects with varying salience or response demands like pedestrians crossing and distracting your way would allow for in-depth understanding of how dynamic elements within the real-life environment impact the cognitive processing and attention allocation

## 5. Summary and Conclusion

In summary, our study weaves a coherent theoretical throughline that connects the chain of exposure, reception, and retention in communication. For messages to have an effect, ensuring exposure to the messages and attention is the key to retention and effects. By bridging the gap between exposure, reception, and retention, we can not only empirically examine the causal influence of theorized variables emotional attention and distraction -, but it also provides a flexible paradigm for future studies on message effects in health communication, politics, or advertising.

## Data Availability Statement

This study’s data are available on Github, at https://github.com/nomcomm/vr_billboard_e.

## References

1. Bettinghaus, E. P. (1986). Health promotion and the knowledge-attitude-behavior continuum. Preventive medicine, 15(5), 475–491. 10.1016/0091-7435(86)90025-3

2. Hornik, R. C. (2002). Exposure: Theory and evidence about all the ways it matters. Social Marketing Quarterly, 8(3), 31–37. 10.1080/15245000214135

3. Reeves, B., Robinson, T., & Ram, N. (2020). Time for the human screenome project. Nature, 577(7790), 314–317. 10.1038/d41586-020-00032-5

4. McGuire, W. J. (1968). Personality and attitude change: An information-processing theory. Psychological foundations of attitudes, 171–196. 10.1016/b978-1-48323071-9.50013-1

5. Potter, W. J. (2008). The importance of considering exposure states when designing survey research studies. Communication Methods and Measures, 2(1-2), 152–166. 10.1080/19312450802062299

6. Broadbent, D (1958). Perception and Communication. London: Pergamon Press. 10.1037/10037-000

7. Lang, A., Bradley, S. D., Park, B., Shin, M., & Chung, Y. (2006). Parsing the resource pie: Using STRTs to measure attention to mediated messages. Media psychology, 8(4), 369–394. 10.1207/s1532785xmep08043

8. James, W. (1890). The principles of psychology (Vol. 1). Cosimo, Inc.

9. Chun, M. M., Golomb, J. D., & Turk-Browne, N. B. (2011). A taxonomy of external and internal attention. Annual Review of Psychology, 62, 73–101. 10.1146/annurev.psych.093008.100427

10. Styles, E. (2006). The psychology of attention. Psychology Press. 10.4324/9780203968215

11. Geisler, W. S., & Cormack, L. K. (2011). Models of overt attention. The Oxford handbook of eye movements, 439–454. 10.1093/oxfordhb/9780199539789.013.0024

12. Driver, J. (2001). A selective review of selective attention research from the past century. British Journal of Psychology, 92(1), 53–78. 10.1348/000712601162103

13. Greenwald, A.G. & Leavitt, C. (1984). Audience involvement in advertising: Four levels. Journal of Consumer Research, 11(1), 581–592. 10.1086/2089941.

14. Chaiken, S. (1980). Heuristic versus systematic information processing and the use of source versus message cues in persuasion. Journal of personality and social psychology, 39(5). 10.1037//0022-3514.39.5.752

15. Kranzler, E. C., Schmälzle, R., O’Donnell, M. B., Pei, R., & Falk, E. B. (2019). Message-elicited brain response moderates the relationship between opportunities for exposure to anti-smoking messages and message recall. Journal of Communication, 69(5), 589–611. 10.1093/joc/jqz035

16. Chun, M. M., & Turk-Browne, N. B. (2007). Interactions between attention and memory. Current Opinion in Neurobiology, 17(2), 177–184. 10.1016/j.conb.2007.03.005

17. Todd, R. M., & Manaligod, M. G. (2018). Implicit guidance of attention: The priority state space framework. Cortex, 102, 121–138. 10.1016/j.cortex.2017.08.001

18. Foulsham, T. (2019). Scenes, saliency maps and scanpaths. Eye Movement Research: An Introduction to Its Scientific Foundations and Applications, 197–238. 10.1007/978-3-030-20085-5_6

19. Schupp, H. T., Flaisch, T., Stockburger, J., & Junghöfer, M. (2006). Emotion and attention: event-related brain potential studies. Progress in Brain Research, 156, 31–51. 10.1016/s0079-6123(06)56002-9

20. Calvo, M. G., & Lang, P. J. (2004). Gaze patterns when looking at emotional pictures: Motivationally biased attention. Motivation and Emotion, 28(3), 221–243. 10.1023/b:moem.0000040153.26156.ed

21. Nummenmaa, L., Hyönä, J., & Calvo, M. G. (2006). Eye movement assessment of selective attentional capture by emotional pictures. Emotion, 6(2), 257. 10.1037/1528-3542.6.2.257

22. Most, S. B., Chun, M. M., Widders, D. M., & Zald, D. H. (2005). Attentional rubbernecking: Cognitive control and personality in emotion-induced blindness. Psychonomic Bulletin & review, 12, 654–661. 10.3758/bf03196754

23. Bradley, M. M. (2009). Natural selective attention: Orienting and emotion. Psychophysiology, 46(1), 1–11. 10.1111/j.1469-8986.2008.00702.x

24. Yiend, J. (2010). The effects of emotion on attention: A review of attentional processing of emotional information. Cognition and Emotion, 221–285. 10.1080/02699930903205698

25. Lang, A. (2000). The limited capacity model of mediated message processing. Journal of communication, 50(1), 46–70. 10.1111/j.1460-2466.2000.tb02833.x

26. Fisher, J.T., Hopp, F.R., & Weber, R. (2023). Mapping attention across multiple media tasks. Media Psychology, 26(5), 505–529. 10.1080/15213269.2022.2161576

27. Simons, D. J. (2000). Attentional capture and inattentional blindness. Trends in Cognitive Sciences, 4(4), 147–155. 10.1016/s1364-6613(00)01455-8

28. Lavie, N. (2005). Distracted and confused?: Selective attention under load. Trends in Cognitive Sciences, 9(2), 75–82. 10.1016/j.tics.2004.12.004

29. Metz, B., Schömig, N., & Krüger, H. P. (2011). Attention during visual secondary tasks in driving: Adaptation to the demands of the driving task. Transportation research part F: traffic Psychology and Behaviour, 14(5), 369–380. 10.1016/j.trf.2011.04.004

30. McGuire, W. J. (1972). Attitude change: The information processing paradigm. Experimental social psychology, 108–141.

31. Craik, F. I., & Lockhart, R. S. (1972). Levels of processing: A framework for memory research. Journal of Verbal learning and Verbal Behavior, 11(6), 671–684. 10.1016/s0022-5371(72)80001-x

32. Kuhl, B. A., & Chun, M. (2014). Memory and attention. In: Anna C. (Kia) Nobre (ed.), Sabine Kastner (ed.). The Oxford Handbook of Attention. OUP. 10.1093/oxfordhb/9780199675111.013.034

33. Cohen, G., & Conway, M. A. (2007). Memory in the real world. Psychology press. 10.4324/9780203934852

34. Potter, R. F., & Choi, J. (2006). The effects of auditory structural complexity on attitudes, attention, arousal, and memory. Media Psychology, 8(4), 395–419. 10.1207/s1532785xmep08044

35. Slater, M. D. (2004). Operationalizing and analyzing exposure: The foundation of media effects research. Journalism & Mass Communication Quarterly, 81(1), 168–183. 10.1177/107769900408100112

36. Southwell, B. G., Barmada, C. H., Hornik, R. C., & Maklan, D. M. (2002). Can we measure encoded exposure? validation evidence from a national campaign. Journal of Health Communication, 7(5), 445–453. 10.1080/10810730290001800

37. Duchowski, A. T. (2017). Eye tracking methodology: Theory and practice. Springer.

38. Miller, L. C., Shaikh, S. J., Jeong, D. C., Wang, L., Gillig, T. K., Godoy, C. G.,… & Read, S. J. (2019). Causal inference in generalizable environments: Systematic representative design. Psychological Inquiry, 30(4), 173–202. 10.1080/1047840x.2019.1693866

39. Parsons, T. D. (2015). Virtual reality for enhanced ecological validity and experimental control in the clinical, affective and social neurosciences. Frontiers in Human Neuroscience, 9, 660. 10.3389/fnhum.2015.00660

40. Platt, J. R. (1964). Strong Inference: Certain systematic methods of scientific thinking may produce much more rapid progress than others. Science, 146(3642), 347–353. 10.1126/science.146.3642.347

41. Spencer, S. J., Zanna, M. P., & Fong, G. T. (2005). Establishing a causal chain: Why experiments are often more effective than mediational analyses in examining psychological processes. Journal of Personality and Social Psychology, 89(6), 845. 10.1037/0022-3514.89.6.845

42. Bonneterre, S. (2023). Evaluating smoking prevention campaigns in virtual reality: an experimental ecological approach (Doctoral dissertation, Université de Nanterre-Paris X).

43. Bonneterre, S., Zerhouni, O., & Boffo, M. (2024). The Influence of Billboard-Based Tobacco Prevention Posters on Memorization, Attitudes, and Craving: Immersive Virtual Reality Study. Journal of Medical Internet Research, 26, e49344.

44. Schmälzle, R., Lim, S., Cho, H. J., Wu, J., & Bente, G. (2023). Examining the exposure-reception-retention link in realistic communication environments via VR and eye-tracking: The VR billboard paradigm. Plos one, 18(11), e0291924. 10.1371/journal.pone.0291924

45. Baayen, R. H., Davidson, D. J., and Bates, D. M. (2008). Mixed-effects modeling with crossed random effects for subjects and items. Journal of Memory and Language. 59, 390–412. 10.1016/j.jml.2007.12.005

46. Clark, H. H. (1973). The language-as-fixed-effect fallacy: A critique of language statistics in psychological research. Journal of Verbal Learning and Verbal Behavior, 12(4), 335–359. 10.1016/s0022-5371(73)80014-3

47. Jackson, S., & Jacobs, S. (1983). Generalizing about messages: Suggestions for design and analysis of experiments. Human Communication Research, 9(2), 169–191. 10.1111/j.1468-2958.1983.tb00691.x

48. Hartmann, T., Wirth, W., Schramm, H., Klimmt, C., Vorderer, P., Gysbers, A.,… & Sacau, A. M. (2016). The spatial presence experience scale (SPES). Journal of Media Psychology, 28(1), 1–15. 10.1027/1864-1105/a000137

49. Kim, H. K., Park, J., Choi, Y., & Choe, M. (2018). Virtual reality sickness questionnaire (VRSQ): Motion sickness measurement index in a virtual reality environment. Applied Ergonomics, 69, 66–73. 10.1016/j.apergo.2017.12.016

50. Johnson, J. D. (2013). Dosage: A guiding principle for health communicators. jRowman & Littlefield.

51. Farley, J., Risko E. F., & Kingstone, A. (2013). Everyday attention and lecture retention: the effects of time, fidgeting, and mind wandering. Frontiers in Psychology, 4, 619. 10.3389/fpsyg.2013.00619

52. Loftus, G. R. (1972). Eye fixations and recognition memory for pictures. Cognitive Psychology, 3(4), 525–551. 10.1016/0010-0285(72)90021-7

53. Neisser, U. (1964). Visual search. Scientific American, 210(6), 94–103. 10.1038/scientificamerican0664-94

54. De Vreese, C. H., & Neijens, P. (2016). Measuring media exposure in a changing communications environment. Communication Methods and Measures, 10(2-3), 69–80. 10.1080/19312458.2016.1150441

55. Nisbett, R. E., & Wilson, T. D. (1977). Telling more than we can know: Verbal reports on mental processes. Psychological Review, 84(3), 231. 10.1037//0033-295x.84.3.231

56. Kingstone, A., Smilek, D., Ristic, J., Kelland Friesen, C., & Eastwood, J. D. (2003). Attention, researchers! It is time to take a look at the real world. Current Directions in Psychological Science, 12(5), 176–180. 10.1111/1467-8721.01255

57. Greenwald, A. G. (2012). There is nothing so theoretical as a good method. Perspectives on psychological science, 7(2), 99–108. 10.1177/1745691611434210

58. Schmälzle, R., & Huskey, R. (2023). Integrating media content analysis, reception analysis, and media effects studies. Frontiers in Neuroscience, 17, 1155750. 10.3389/fnins.2023.1155750

59. McGuire W.J. (2001). Input and Output Variables Currently Promising for Constructing Persuasive Communications. In: Rice R.E., Atkin C.K., editors. Public Communication Campaigns. SAGE Publications, Inc.; Thousand Oaks, CA, USA. pp. 22–48. 10.4135/9781452233260.n2

60. Cho, H. (Ed.). (2011). Health communication message design: Theory and practice. Sage Publications.

61. Werner, J. S., & Chalupa, L. M. (Eds.). (2013). The new visual neurosciences. MIT Press.

62. Turner, M. M. (2012). Using emotional appeals in health messages. In Cho, H. Health communication message design: Theory and practice, 59–72.

63. Mongeau, P. A. (2013). Fear appeals. The SAGE handbook of persuasion: Developments in theory and practice, 184–199. 10.4135/9781452218410.n12

64. Janis, I. L., & Feshbach, S. (1953). Effects of fear-arousing communications. Journal of Abnormal and Social Psychology, 48(1), 78. 10.1037/h0060732

65. Ihssen, N., & Keil, A. (2009). The costs and benefits of processing emotional stimuli during rapid serial visual presentation. Cognition and Emotion, 23(2), 296–326. 10.1080/02699930801987504

66. Schupp, H. T., Kirmse, U., Schmälzle, R., Flaisch, T., & Renner, B. (2016). Newly-formed emotional memories guide selective attention processes: Evidence from event-related potentials. Scientific Reports, 6, 28091. 10.1038/srep28091

67. Prentice, D. A., & Miller, D. T. (1992). When small effects are impressive. Psychological Bulletin, 112(1), 160–164. 10.1037/0033-2909.112.1.160

68. Nabi, R. L. (1999). A cognitive functional model for the effects of discrete negative emotions on information processing, attitude change and recall. Communication Theory, 9, 292–320. 10.1111/j.1468-2885.1999.tb00172.x

69. O’Keefe, D. J., & Hoeken, H. (2021). Message design choices don’t make much difference to persuasiveness and can’t be counted on—Not even when moderating conditions are specified. Frontiers in Psychology, 12, 664160. 10.3389/fpsyg.2021.664160

70. Turner, M. M., Skubisz, C., Pandya, S. P., Silverman, M., & Austin, L. L. (2014). Predicting visual attention to nutrition information on food products: the influence of motivation and ability. Journal of health communication, 19(9), 1017–1029. 10.1080/10810730.2013.864726

71. Li, K., Huang, G., & Bente, G. (2016). The impacts of banner format and animation speed on banner effectiveness: Evidence from eye movements. Computers in Human Behavior, 54, 522–530. 10.1016/j.chb.2015.08.056

72. Yzer, M., Han, J., & Choi, K. (2018). Eye movement patterns in response to anti-binge drinking messages. Health communication, 33(12), 1454–1461. 10.1080/10410236.2017.1359032

73. Nastase, S. A., Goldstein, A., & Hasson, U. (2020). Keep it real: rethinking the primacy of experimental control in cognitive neuroscience. NeuroImage, 222, 117254. 10.1016/j.neuroimage.2020.117254

74. Farahany, N. A. (2023). The battle for your brain: defending the right to think freely in the age of neurotechnology. St. Martin’s Press.

